# A sweeter future: Using protein language models for exploring sweeter brazzein homologs

**DOI:** 10.1101/2023.02.20.529172

**Authors:** Bryan Nicholas Chua, Wei Mei Guo, Han Teng Wong, Dave Siak-Wei Ow, Pooi Leng Ho, Winston Koh, Ann Koay, Fong Tian Wong

## Abstract

Reducing sugar intake lowers the risk of obesity and associated metabolic disorders. Currently, this is achieved using artificial non-nutritive sweeteners, where their safety is widely debated and their contributions in various diseases is controversial. Emerging research suggests that these sweeteners may even increase the risk of cancer and cardiovascular problems, and some people experience gastrointestinal issues as a result of using them. A safer alternative to artificial sweeteners could be sweet-tasting proteins, such as brazzein, which do not appear to have any adverse health effects.

In this study, protein language models were explored as a new method for protein design of brazzein. This innovative approach resulted in the identification of unexpected mutations, which opened up new possibilities for engineering thermostable and potentially sweeter versions of brazzein. To facilitate the characterization of the brazzein mutants, a simplified procedure was developed for expressing and analyzing related proteins. This process involved an efficient purification method using *Lactococcus lactis* (*L. lactis*), a generally recognized as safe (GRAS) bacterium, as well as taste receptor assays to evaluate sweetness. The study successfully demonstrated the potential of computational design in producing a more heat-resistant and potentially more palatable brazzein variant, V23.

## 1. Introduction

The dramatic rise in obesity and diabetes in recent decades has seen the widespread use of artificial sweeteners in food and drinks as sugar replacements (Gardner et al., 2012). By replacing sugar with these artificial sweeteners, blood glucose level can be better regulated and calorie consumption reduced whilst maintaining food palatability with its sweet taste (Gardner et al., 2012). However, recent data around the detrimental side effects of consuming artificial sweeteners highlight the need for other sweeteners (Bueno-Hernández et al., 2019; Debras, Chazelas, Sellem, et al., 2022; Debras, Chazelas, Srour, et al., 2022; Suez et al., 2014).

Sweet proteins of natural origin have the potential to replace these artificial sweeteners due to their intensely sweet nature and low risk safety profile (Gu et al., 2015; Rega et al., 2015). Unlike sucrose, sweet proteins do not trigger a demand for insulin in diabetic patients, (Gnanavel & Serva Peddha, 2011; Gu et al., 2015). So far, seven vastly different sweet-tasting proteins have been discovered from plants located in tropical rainforests. These are brazzein, thaumatin, monelin, neoculin, mabinlin, miraculin, and pentadin (Zhao et al., 2021). Sweet proteins bring about sweet taste perception in humans by interacting with the human sweet taste receptor TAS1R2/TAS1R3 (Kim et al., 2022). Of the sweet proteins, the most promising candidate under consideration for direct sugar replacement is brazzein, due to its relatively small size of 54 amino acids with 6.40 kDa (Caldwell et al., 1998), and its intense sweetness that is 500 to 2000 times over sucrose. (Assadi-Porter, Aceti, Cheng, et al., 2000). Originally isolated from the fruit of the west African plant, *Pentadiplandra brazzeana* Bailon (Ming & Hellekant, 1994), brazzein’s thermal and pH stability make it an ideal system for application in the biotechnology and food industries.

For scalable sustainable food production, the food industry has also initiated the implementation of precise fermentation techniques in the creation of protein-based food ingredients. (Teng et al., 2021). These innovative food products offer healthier and more sustainable options for climate-conscious consumers and have the potential to change our understanding of food. Simultaneously, there have been significant advancements in the field of protein engineering, aided by the emergence of computational techniques and algorithms (Marchand et al., 2022; Meinen & Bahl, 2021). This has created new avenues for designing proteins with improved characteristics such as enhanced stability, activity, and specificity, including generation of food ingredients with advanced features such as improved taste, texture, and nutritional value.

In this study, we explored an alternative technique of protein design by using protein language models as hypothesis generators. In our approach, we projected the wild type (WT) protein sequence of brazzein into the embedding space, following which we performed a random walk to explore potentially new and diverse sequences around the vicinity of WT brazzein. These sequences were then sampled, identified and expressed. This exploration resulted in brazzein sequences with unexpected mutations, offering new unconventional starting points for engineering thermostable variations with a potential for increased sweetness. Furthermore, to accelerate characterisation and to move towards a viable production strategy as a food ingredient, we have also developed a faster and more efficient protocol for purifying these brazzein variants from generally recognized as safe (GRAS) bacteria *Lactococcus lactis* (*L. lactis*).

## 2. Materials & Methods

### 2.1. Plasmid construction

*Escherichia coli* (*E*. c*oli*) codon-optimized DNA sequences for the His-tagged brazzein constructs were synthesized in a pET24a(+) vector from Twist Biosciences. The constructs were transformed into *E. coli* OmniMAX™2 for sequencing and into BL21(DE3) *E. coli* for protein expression. *L. lactis* codon-optimized DNA sequences for the His-tagged brazzein constructs were synthesized from Twist Biosciences. The fragments were cloned into pNZ8148 vector via Gibson assembly. The constructs were transformed into *L. lactis* NZ9000 for sequencing and protein expression.

### 2.2. BL21(DE3) *E. coli* protein expression and purification

Single colonies from transformed BL21(DE3) *E. coli* were inoculated in Luria-Bertani (LB) broth containing 50 μg/mL kanamycin for overnight culture at 37 °C, with shaking at 200 rpm. The overnight cultures were inoculated into 300 mL Terrific Broth containing 50 μg/mL kanamycin at 37 °C, 200 rpm. When the optical density at 600nm (OD_600_) reached 0.4 – 0.6, protein expression was induced with 1 mM of isopropyl β-D-1-thiogalactopyranoside (IPTG), and incubated overnight at 30 °C, 200 rpm. The cultures were subsequently harvested by centrifugation at 8000 *g* for 10 min at 4 °C. The resulting pellets were freeze-thawed, resuspended in BugBuster Protein Extraction Reagent (Merck, Cat. No. 70584) as per vendor’s instruction and then incubated at room temperature for 15 min with rotation. The resulting lysate was then centrifuged at 18000 *g* for 20 min at 4 °C. The supernatant was incubated with PureCube 100 INDIGO Ni-Agarose resin (Cube-biotech, Cat. No. 75110) for 1 h at room temperature, and the protein-bound resin was washed with 50 mM sodium phosphate buffer pH 7.4, 500 mM sodium chloride and 20 mM imidazole. The bound protein was eluted with 50 mM sodium phosphate buffer pH 7.4, 500 mM sodium chloride and 500 mM imidazole. The eluate was buffer-exchanged and concentrated with Hank’s Balanced Salt Solution (HBSS) containing 20 mM 4-(2-hydroxyethyl)-1-piperazineethanesulfonic acid (HEPES) at pH 7.0 using a spin concentrator with 3kDa MWCO.

### 2.3. *L. lactis* NZ9000 protein expression and purification

Single colonies from the transformed *L. lactis* NZ9000 were inoculated in M17 Broth (0.5 % glucose,10 μg/mL chloramphenicol) and incubated overnight at 30 °C without shaking. The overnight cultures were inoculated in 2 L 2x M17 Broth (2 % glucose, 10 μg/mL chloramphenicol) to OD_600_ 0.1, and incubated at 30 °C. When OD_600_ reached 1.0, protein expression was induced with 50 ng/mL nisin for 3 h at 30 °C.

The cultures were centrifuged at 8000 *g* for 10 min at 4 °C. The resulting pellets were freeze-thawed, resuspended in 50 mM sodium phosphate buffer pH 7.4, 300 mM sodium chloride, 10 mM imidazole and 0.03 % Triton X-100, and then incubated at room temperature for 15 min. The resuspended pellets were either sonicated 4 times for 10 s at 10 s intervals on ice or heated at 95 °C for 10 min. The resulting lysate was then centrifuged at 18000 *g* for 20 min at 4 °C. The protein was purified from the supernatant as described earlier. For thermostability testing (Fig 3), the samples in HBSS-HEPES buffer were heated at 95 °C for 4 h then rapidly cooled to 4 °C.

### 2.4. Sweet taste receptor luminescence assay

HEK 293T (ATCC) cells were seeded at a density of 20,000 cells per well in white 384-well tissue culture plates (Greiner), coated with Poly-D-Lysine (PDL; Sigma) at a final concentration of 1 mg/mL. After an overnight incubation at 37 °C in a humidified atmosphere of 5 % CO2, the cells were transiently transfected with two plasmids. The first plasmid is a multigene CMV-promoter based expression vector containing the genes for the sweet taste receptor (TAS1R2/TAS1R3) and the chimeric Gα16-gust44 gene. The second plasmid expression vector contains the gene for the apophotoprotein, mitochondrial-targeted (mt)-clytin II. The two plasmids were transfected at a ratio of 20 ng:20 ng per well using ViaFect (Promega), employing a transfection agent to plasmid ratio of 3:1 μL:μg.

At 6 h post-transfection, the culture media was replaced with low-glucose DMEM (Gibco) supplemented with 10 % (v/v) dialysed FBS (Biowest) and 1 % (v/v) penicillin-streptomycin (Gibco). The following day, the spent media in the wells with transfected cells were removed, leaving a residual volume of 10 μL/well. The cells were loaded with additional 25 μL of coelenterazine F (AAT Bioquest) to a final concentration of 10 μM, in assay buffer (1× HBSS assay buffer with 20 mM HEPES at pH 7.0) and incubated for 4 hours at 27 °C in the dark.

The assay was performed using the luminescence mode of the Fluorescent Imaging Plate Reader (FLIPRTETRA, Molecular Devices) controlled by the ScreenWorks software (version 4.0.0.30, Molecular Devices). A baseline read was captured for an initial 10 s before 25 μL of test ligand prepared to a 2.4 times concentration in assay buffer was dispensed from the source plate into the assay plate. The kinetic data was acquired for a further 100 s to record the responses of each well to added test sample. The well responses were exported as area under the curve (AUC) values and the data were plotted using the four-parameter logarithmic regression equation using Prism 8 (GraphPad) software. The data reported were derived from at least two independent experiments, performed in duplicates. For this study, we used two reference sweeteners, sucralose, and the sweet protein thaumatin for comparison to our brazzein test samples (Joseph et al., 2019).

### 2.5. Sweet taste receptor fluorescence assay

293AD (Cell Biolabs, Inc) cells were maintained under similar cell culture conditions as 293T cells. Cells were seeded to a density of 12,000 cells per well in black 384-well tissue culture plates (Greiner) and grown overnight. A multigene CMV-promoter based expression vector containing the genes for TAS1R2/TAS1R3 and chimeric Gα16-gust44 was transiently transfected into 293AD cells at 25 ng per well using Viafect reagent. After 6 h post-transfection, the growth media was removed and replaced with low-glucose DMEM (Gibco) supplemented with 10 % (v/v) dialysed FBS (Biowest) and 1 % (v/v) penicillin-streptomycin (Gibco). The following day, the transfected cells were loaded with Calcium 6 (Molecular Devices) fluorescent dye. The assay plate was first incubated at 37 °C in a humidified incubator with 5 % CO_*2*_ for 2 h, followed by a further 30 min on the lab bench for equilibration to ambient temperature.

The assay was performed using the fluorescence mode of the FLIPR-TETRA. The fluorescence intensity is directly correlated to the amount of intracellular calcium that is released into the cytoplasm in response to ligand-mediated activation of the sweet taste receptor, which in turn is regarded as a measure of receptor activation. Changes in calcium membrane potential were measured over time by fluorescence measurements with an excitation at 470−495 nm and measurement of emission at 515−575 nm. A baseline measurement read was taken every second for 10 s prior to addition of sweetener or test sample, where further measurement reads were acquired for 310 s.

Emission fluorescence values were converted to response over baseline values using the ScreenWorks software (version 4.0.0.30, Molecular Devices), and the data was plotted using the four-parameter logarithmic regression equation using Prism 8 (GraphPad) software. Positive assay responses by samples can be evaluated for its potency towards the sweet taste receptor, expressed as EC_50_, which is the concentration of molecule required to a give half-maximal response in the sweet taste receptor assay.

### 2.6. Statistical tests

With the comparison of protein yields from 2 lysis methods, data is presented as mean with standard deviation (SD) from 3 replicates over 3 independent runs. Unpaired t-test of protein yields between 2 lysis methods was performed using GraphPad Prism to determine statistical significance. With brazzein samples produced in *L. lactis* and tested in the fluorescence-based sweet taste receptor assay, collected data was fitted in GraphPad Prism using a four-parameter logistic fit equation. Data interpolation was performed at a single protein concentration of 68 μg/mL and presented as mean with SD from at least four experimental replicates. One-way analysis of variance (ANOVA) and Tukey’s multiple comparison test were employed to compare data. P values less than 0.05 were considered statistically significant.

## 3. Results & Discussion

### 3.1. Generation of computationally fold-able sweet protein variants

Protein design has been traditionally approached through rational protein engineering and ancestral sequence reconstruction. In ancestral reconstruction, the ancestral sequences are inferred from related protein sequences and then modified to attain desired properties. In rational protein engineering, the desired properties are optimized by introducing mutations based on experimental data on how they impact protein structure. Both of these approaches rely heavily on sequences that have already been optimized by nature as starting points. In our work, we are interested in exploring beyond nature’s optimized set of sequences. To achieve this, we employ protein language models to explore new starting points for protein engineering. Pretrained protein language models (SeqVec (Heinzinger et al., 2019), UniRep (Alley et al., 2019), CPCprot (Lu et al., 2020)) are capable of learning a generalized description of proteins by representing protein sequences as a language and converting them into numerical representations in a multidimensional latent space that encompasses all other possible protein sequences. Prior studies (Alley et al., 2019; Heinzinger et al., 2019; Lu et al., 2020) have shown that the latent space close to the protein of interest contains sequences that preserve both the structural and functional properties of the original protein. Leveraging this property and using WT brazzein sequence as the initial starting point, we introduced multiple mutations to sample sequences in the latent space that surrounds WT brazzein. The sampled sequences are then presented to human scientists for guidance and rational curation of desired properties such as: 1) novelty & divergence in sequence, 2) thermostability and 3) solubility. This human directed process of evolution of the sequence along different dimensions in the latent space allowed us to construct a library of sequences as potential leads for downstream characterization (Fig 1).

**Figure 1.**
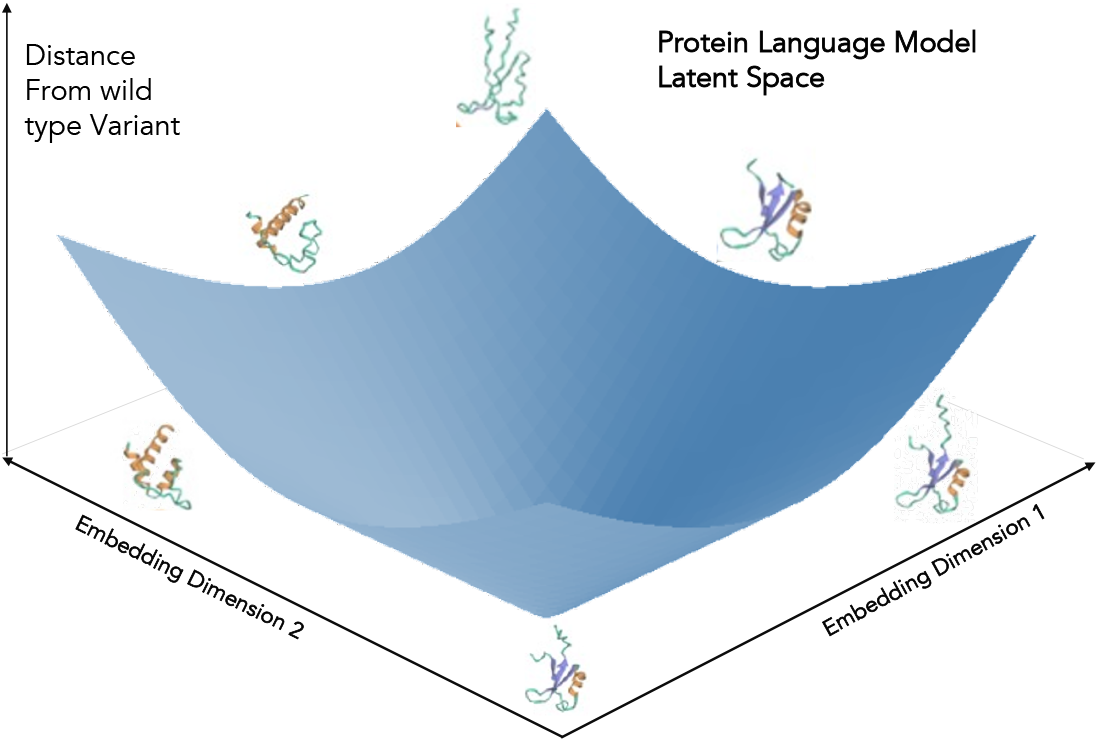
Overall strategy of designing proteins by exploring the latent space of Large Language Models trained on protein sequences. The process of generating a library of variants begins with the selection of an initial native protein with the desired sweetness profile.

### 3.2. Scaling up throughput of sweetness measurement and comparison

In this study, we employed the use of cell-based sweet taste receptor assays to assess the relative sweetness of brazzein variants in a rapid and systematic manner. Similar receptor-based assays have been routinely used in studies of sweet taste reception and sweetener molecules optimization (Riedel et al., 2017). Human sweet taste receptors, along with their signaling components are heterologously expressed in cultured mammalian HEK cells and shown to respond to a wide array of sweeteners. This approach measures calcium mobilization in response to sweet taste receptor activation by sweeteners such as sweet proteins like brazzein, carbohydrate sweeteners like sucrose, and both natural and synthetic sweetener molecules such as sucralose and stevioside. This allows rapid and increased throughput of screening which otherwise would not be possible with a human sensory panel.

An initial library of five brazzein variants, V21-V25, were generated computationally, with a range of 5-8 mutations, including deletions (Fig 2A). As an initial assay, the variants were expressed in *E. coli* and purified via affinity tag pull-down before screening for sweet taste receptor response and compared against WT brazzein as a control. As no optimization for purification of brazzein was performed in this initial set of screening experiments, the samples expressed in *E. coli* had a high percentage of impurity (Fig S1), which contributed to non-specific signals in our assay readout. Consequently, we cannot wholly attribute the observed assay signal specifically to the sweet taste receptor activation, so we are only able to have approximate relative sweetness. This initial variant library showed a trend where the V23 variant could potentially be sweeter than WT (Fig 2B). To further investigate this, we focused on producing high purity samples of the V23 variant alongside WT brazzein.

**Figure 2.**
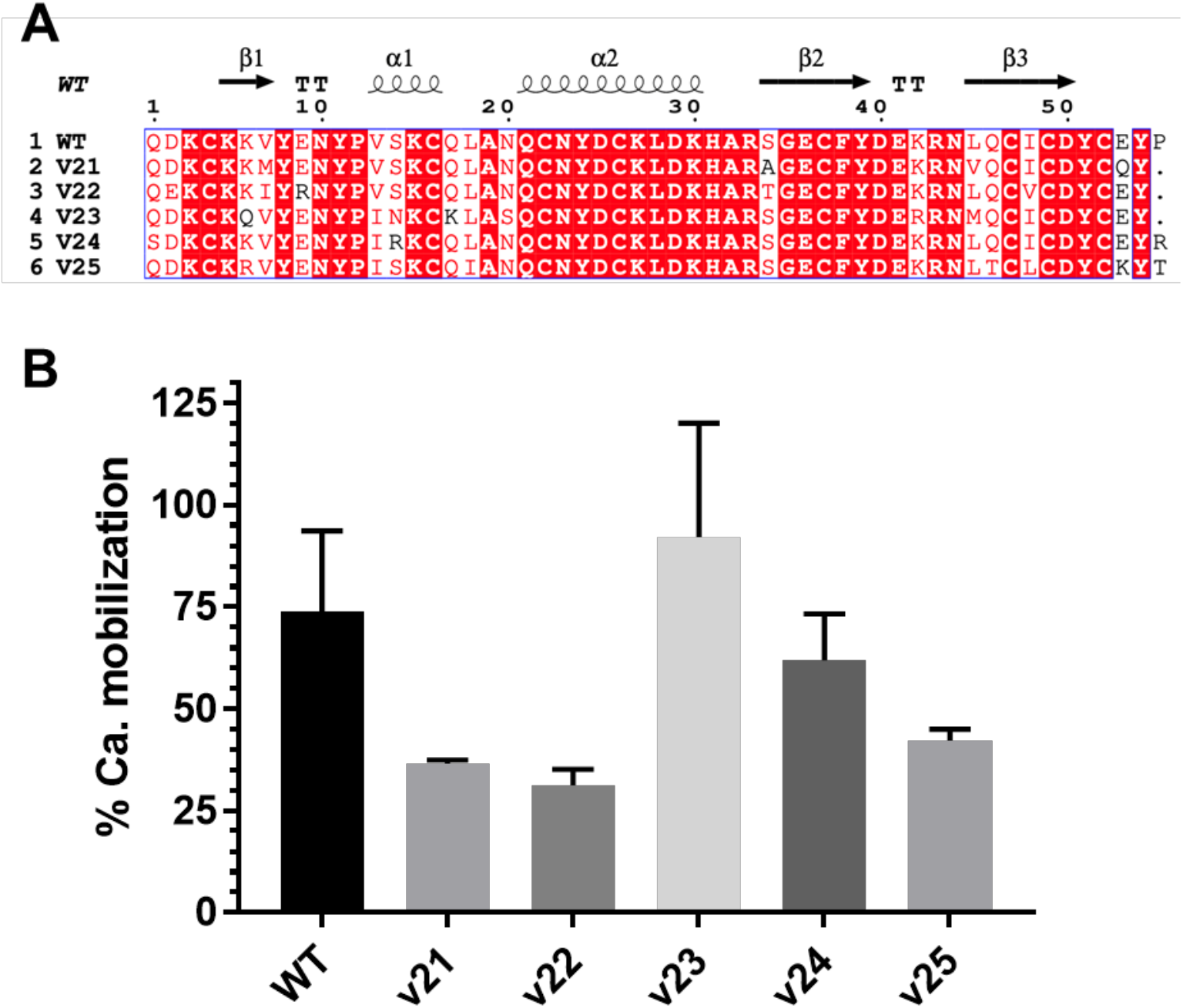
Artificial intelligence (AI)-derived mutants. (A) Sequence alignment of mutants (V21-25) against WT brazzein. Identical residues are highlighted in red. Similar residues are colored red and residues that are not similar are colored black. Structural elements of WT brazzein are also shown (PDB: 4HE7). The numbering starts from Q after the N-terminal M. (B) Comparison of sweet taste receptor responses using calcium mobilization responses of various mutants and WT brazzein expressed in *E. coli* using the luminescence-based readout sweet taste receptor assay. Percent calcium mobilization is calculated against the maximum assay response from 15 mM sucralose. Data is interpolated and averaged from non-linear fits of experimentally derived data of at least two independent assay runs at a standardized protein concentration of 0.1 mg/mL. Error bars are SD.

### 3.3. Bioprocessing optimization in a *L. lactis*

To optimize expression of brazzein for characterisation, we utilized GRAS *L. lactis* NZ9000 (Berlec & Strukelj, 2009; Linares et al., 2010). Although brazzein purified from *L. lactis* had high purity (Fig. 3), its yields are significantly low; < 0.1 mg/L. Subsequently, we also exploited brazzein’s thermostability (Ming & Hellekant, 1994) to establish a heat-based purification protocol. While both heat lysis and sonication effectively rupture bacteria cells to release intracellular proteins, heat lysis is particularly effective when working with heat-stable proteins (Kalthoff, 2003). Others have previously used a 2 h heat treatment at 80 °C as a second purification step after ammonium precipitation to successfully increase the purity of brazzein expressed in transgenic tobacco leaves (Choi et al., 2022). Hence, we hypothesize that heating can be used in lieu of mechanical lysis (sonication) to lyse and purify the expressed brazzein. Heat-based lysis of cell pellets was performed by heating cell pellets at 95 °C for 10 min (Fig 3A). In our observations, purity of the samples increased with heat lysis protocol (Fig 3B). More importantly, there was also a significant 10-fold increase in brazzein yield between the two protocols (Fig 3C). We reason that the improvement in purity and yield could be due to the removal of all non-heat-stable proteins through denaturation, including heat denaturation of proteases that could otherwise break down the target protein (Kalthoff, 2003). Although we did not explore further in this study, we expect that yields can be further improved through an in-depth optimization study of fermentation conditions combined with heat lysis-mediated purification (Berlec et al., 2008).

### 3.4. Characterization of AI-derived V23 variant

Using *L. lactis* as our expression host, we examined the sweetness response of sonicated and heat lysed products of WT brazzein and V23. Thermostability was also further tested by heating the brazzein samples for a further 4 h at 95 °C (Fig 3A – treatment 3). In comparison with WT’s sonicated and heat treated products, V23 equivalents had higher responses and potencies in the sweet taste receptor assay (Fig 4, Table S1). This suggests that V23 is potentially sweeter than WT brazzein and maintains this potency with heat treatment. Overall, heat treated products are less sensitive in the taste receptor assay compared to sonicated products, which is not surprising since we expect some protein folding to be disrupted in the presence of heat. Interestingly, the samples that were heated for 4 h had slightly higher calcium response compared to 10 min lysis, suggesting that further heating might have removed residual heat labile proteins including slightly misfolded brazzein proteins. Even though there seems to be a drop in sweet potency of products from 10 min heat lysis and sonication protocols, prior examples of heat treatment protocols at mild conditions, for example, 80 °C, 2 h (Choi et al., 2022), imply that further refinement of the heat treatment could be utilized to minimize impact on sweetness potency while preserving yields.

**Figure 3.**
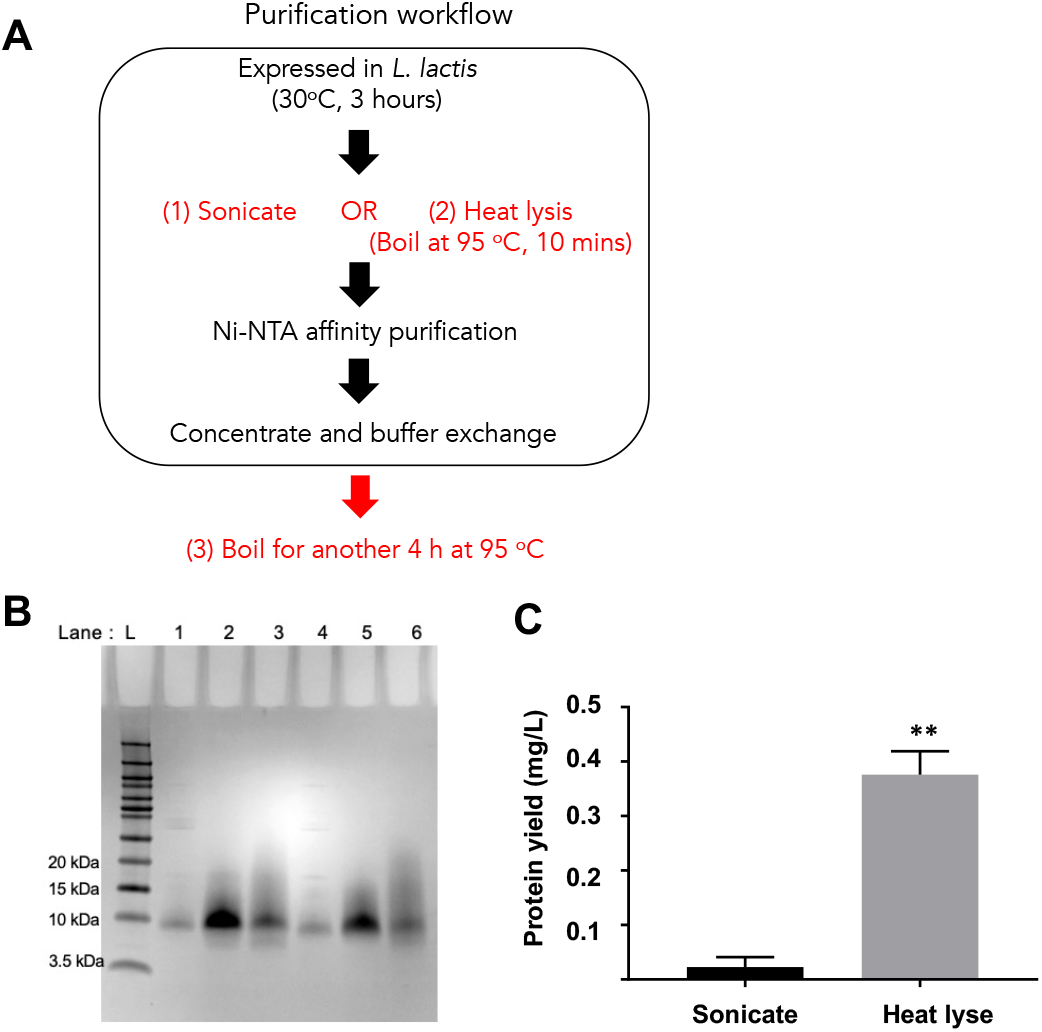
Expression and processing of brazzein from GRAS *L. lactis*. (A) Purification workflow (B) Representative protein gel of purified WT brazzein with L: Novex pre-stained ladder, lane 1: sonicated (Fig 3A – treatment 1), lane 2: heat lysis (95 °C for 10 min, Fig 3A – treatment 2), lane 3: heat lysis, purified and heated 4 h at 95 °C (Fig 3A – treatment 3) and its equivalent for V23 (lanes 4-6). Invitrogen Novex 16 % Tricine gel was used with Tricine SDS Running buffer. The gel was stained with Coomassie blue and imaged. (C) Total determined protein concentration of His-tagged purified brazzein with two lysis methods (Fig 3A – treatments 1 and 2). Data are from 3 replicates over 3 independent runs. Error bars are SD. Unpaired t-test of protein yields between two lysis methods reveal statistical significance (*p* < 0.05).

**Figure 4.**
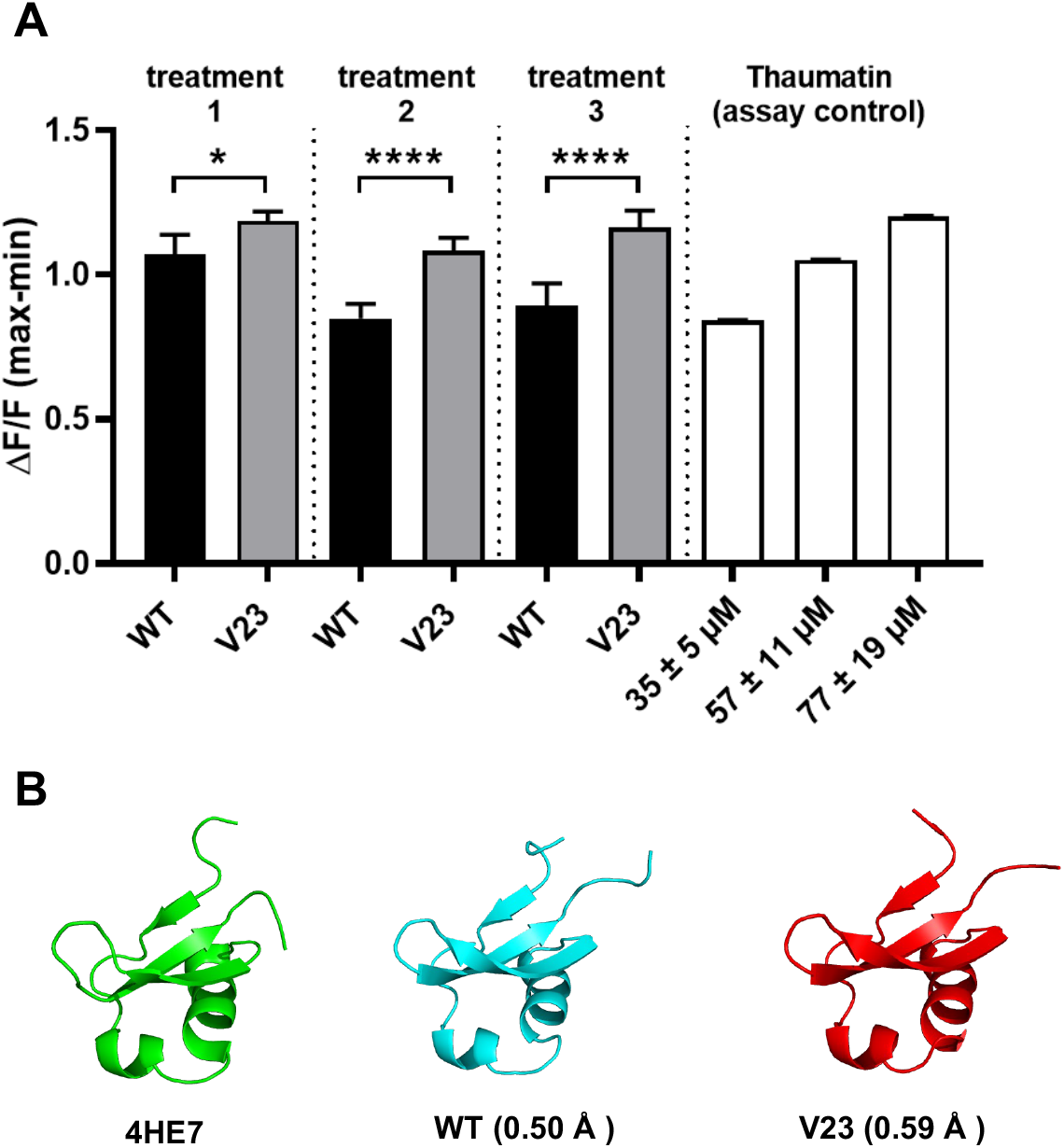
Analysis of V23 (A) Sweet potencies using calcium mobilization responses of WT brazzein and V23, expressed in *Lactococcus lactis* (*L. lactis*) and subjected to three different treatments. Treatment 1: sonicated, no heat treatment, 2: 10 min at 95 °C, 3: 10 min + 4 h at 95 °C (See Fig 3 for more details). Samples were tested using the fluorescence-based sweet taste receptor assay. Thaumatin is shown for scale of sweet taste receptor assay response at various concentrations. Data is interpolated and averaged from non-linear fits of experimentally derived data at a single protein concentration point of 68 μg/mL, from at least six experimental replicates. Error bars are SD. One asterisk (*) indicates *p* = 0.0138 (one-way ANOVA). Four asterisks (****) indicate *p* < 0.0001. (B) Predicted structures were generated and compared using ESMfold (Lin et al., 2022). RMSD of the predicted structures against 4HE7 (experimentally derived structure) is given in brackets. WT brazzein sequence is used as a control for folding.

## 4. Conclusion

In this study, we showed the validity of the hypothesis that novel amino acid sequences generated by protein language models and situated near the target protein in the latent space are capable of retaining both the structural and functional features of the original protein. Additionally, we showcased that exploration of this space can be used to funnel down to highly optimized candidates that would not be obvious using conventional methods. We have also shown that such a library can be use to screen and create thermostable and potentially sweeter brazzein homologs. To characterize these homologs, we also established an efficient and systematic workflow for expression and characterization of brazzein. This includes quality control assays that quantify the sweetness of proteins, enabling accurate characterization and comparison of brazzein mutants, as well as a simplified but productive purification protocol from the GRAS *L. lactis* for consistent production of high-purity brazzein. By combining *in silico and in vitro* workflows, we demonstrated the application of computational design to create a thermostable and potentially sweeter brazzein homolog, V23.

In this study, AI-powered protein design allows us to quickly evaluate a large number of protein sequences which in turn allows us to identify highly optimized candidates that would be hard to uncover with conventional methods. Without *a* prior input of brazzein -specific data, we were able to design high order functional mutants (5-8 mutations), 9-12 % of the total protein length, where a screen of 5 mutants uncovered one better than WT brazzein. Prior studies on WT brazzein have examined single or double mutations to uncover key areas vital for sweetness. The results indicate that the residues 29-33, 36, 39-43, and the N and C-termini are crucial for sweetness (Assadi-Porter, Aceti, & Markley, 2000; Ghanavatian et al., 2016; Jin, Danilova, Assadi-Porter, Aceti, et al., 2003; Jin, Danilova, Assadi-Porter, Markley, et al., 2003; Lee et al., 2013; Lim et al., 2016; Liu et al., 2021; R. Lu et al., 2022). With computationally generated mutations, we were directed to regions different from these previously studied regions. Furthermore, our mutations were not predicted by prior studies focused on predicting the thermostability of brazzein (Tang et al., 2021). This implies that there is still signficiant potential for enhancing the sweetness of brazzein beyond regions that have been explored through traditional protein engineering, and vice versa, the computationally designed variant can be further improved by combining it with already known mutations.

## Abbreviations

AI: Artificial intelligence
ANOVA: Analysis of variance
ATCC: American Type Culture Collection
AUC: Area under curve
CMV: Cytomegalovirus
DMEM: Dulbecco’s Modified Eagle Medium
EC_50_: Concentration of a ligand that leads to 50 % maximal response
*E*. c*oli*: *Escherichia coli*
FBS: Fetal bovine serum
FLIPR: Fluorescence Imaging Plate Reader
GRAS: Generally recognized as safe
HBSS: Hank’s Balanced Salt Solution
HEK: Human embryonic kidney
HEPES: 4-(2-hydroxyethyl)-1-piperazineethanesulfonic acid
IPTG: Isopropyl β-D-1-thiogalactopyranoside
LB: Luria-Bertani
*L.*: *lactis Lactococcus lactis*
MT: Mitochondrial-targeted
PDL: Poly-D-Lysine
SD: Standard deviation
SDS: Sodium Dodecyl Sulfate
TAS: Taste receptor
WT: Wild type

## Acknowledgements

We gratefully acknowledge financial support from the Singapore Food Story R&D Program (W20W2D0011 & H20H8a0003) and Agency for Science, Technology and Research (A*STAR), Singapore (#21719) for this work.

## Data Availability

The data used in this study can be found in the main text and supplementary materials of the paper.

## Author Contributions

Bryan Nicholas Chua: designed and conducted protein expression experiments, analysis, drafted original manuscript and figures.

Wei Mei Guo: designed and conducted assays, analysis, drafted original manuscript and figures.

Han Teng Wong: conceptualization, designed and conducted protein expression experiments, reviewed manuscript

Dave Siak-Wei Ow: designed *L. lactis* experiments, reviewed manuscript Pooi Leng Ho: conducted *L. lactis* experiments, reviewed manuscript

Winston Koh: conceptualization, designed and conducted computational experiments, drafted original manuscript and figures.

Ann Koay: conceptualization, designed and conducted assays, analysis, drafted original manuscript and figures.

Fong Tian Wong: conceptualization, analysis, designed and conducted protein expression experiments, drafted original manuscript and figures.

## Declaration of Competing Interest

A patent application has been filed by the authors.

**Appendix A. Supplementary data**

**Fig S1**. WT brazzein and AI-designed variants of brazzein expressed in *E. coli*.

**Table S1**. EC_50_ values, expressed in μM, of wild type brazzein and AI-designed V23 mutants, subjected to three different treatment methods (see Fig 2).

## Notes

### Summary of Updates

Declaration updated.

