## Supplemental data for "A sweeter future: Using protein language models for exploring sweeter brazzein homologs"

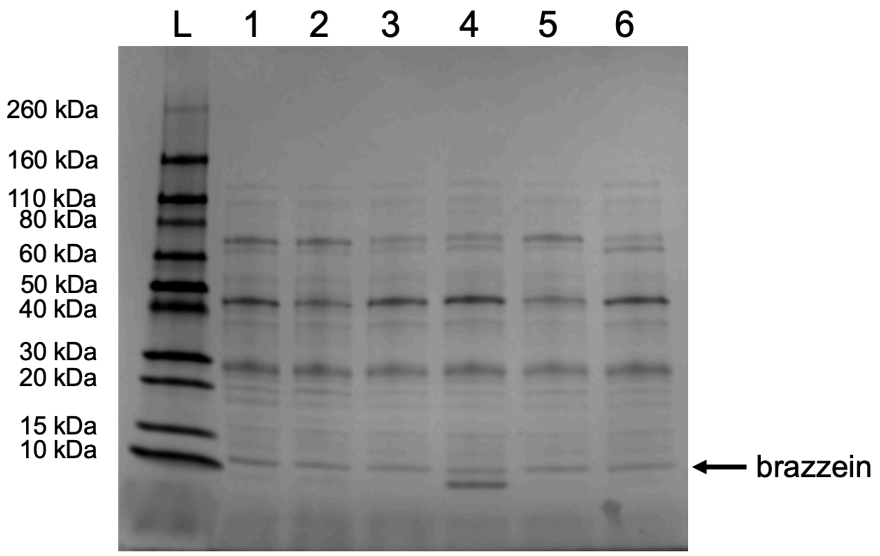


**Figure S1.** Wild type brazzein and AI-designed variants of brazzein were expressed in *E. coli*. Cells were lysed using Bugbuster and the lysate was purified by Ni-NTA affinity chromatography. The purified lysate was concentrated, buffer exchanged and run on a 4-20% Biorad Mini-PROTEAN TGX Stain-Free gel with Tris-Glycine-SDS buffer. The gel was stained with Coomassie blue and imaged. L: Novex pre-stained ladder, 1: brazzein thermal variant 21, 2: brazzein thermal variant 22, 3: brazzein thermal variant 23, 4: brazzein thermal variant 24, 5: brazzein thermal variant 25, and 6: wild-type brazzein.

**Table S1:** $EC_{50}$ values, expressed in µM, of wild type brazzein and AI-designed V23 mutants, subjected to three different treatment methods (see Fig 2). The samples were tested using the fluorescence based assay

| | **Treatment 2** | | **Treatment 3** | | **Treatment 1** | |
|---|---|---|---|---|---|---|
| **$EC_{50}$ (µM)** | **WT** | **V23** | **WT** | **V23** | **WT** | **V23** |
| curve 1 | 15.1 | 10.3 | 25.6 | 3.2 | 1.9 | 1.3 |
| curve 2 | 20.5 | 11.7 | 22.9 | 4.9 | 5.2 | 8.8 |
| curve 3 | 16 | 12.9 | 22.1 | 3.5 | 1.6 | 5.6 |
| curve 4 | 25.6 | 8.3 | 14.7 | 5.3 | 11.9 | 2.1 |
| curve 5 | 17.4 | 8.3 | 11 | 3.1 | 2.4 | 1.7 |
| curve 6 | 23.3 | 12.5 | 12.5 | 5.2 | 10.1 | 1.1 |
| **Mean and S.D.** | **19.6 ± 4.2** | **10.6 ± 2.1** | **18.1 ± 6.2** | **4.2 ± 1.1** | **5.5 ± 4.5** | **3.4 ± 3.1** |

The $EC_{50}$ values were generated using the four-parameter logarithmic regression equation in Prism 8 (GraphPad) software with the following constraints applied: bottom asymptote > 0.25; 0 < top asymptote < 2.
